# Semi-coordinated allelic-bursting shape dynamic random monoallelic expression in pre-gastrulation embryos

**DOI:** 10.1101/2020.09.18.303776

**Authors:** C H Naik, K Hari, D Chandel, S Mandal, M K Jolly, S Gayen

## Abstract

In recent years, allele-specific single-cell RNA-seq (scRNA-seq) analysis has demonstrated wide-spread dynamic random monoallelic expression of autosomal genes (aRME) in different cell types. However, the prevalence of dynamic aRME during pre-gastrulation development remains unknown. Here, we show that dynamic aRME is wide-spread in different lineages of pre-gastrulation embryos. Additionally, the origin of dynamic aRME remains poorly understood. Theoretically, it is believed that independent transcriptional bursting from each allele leads to the dynamic aRME. However, based on analysis of allele-specific burst kinetics of autosomal genes, we found that allelic burst is not perfectly independent, rather it happens in semi-coordinated fashion. Importantly, we show that semi-coordinated allelic bursting of the genes; particularly with low burst frequency, leads to frequent asynchronous allelic bursting, thereby shaping the landscape of dynamic aRME in pre-gastrulation embryos. Altogether, our study provides significant insight into the prevalence and origin of dynamic aRME and cell to cell expression heterogeneity during early mammalian development.

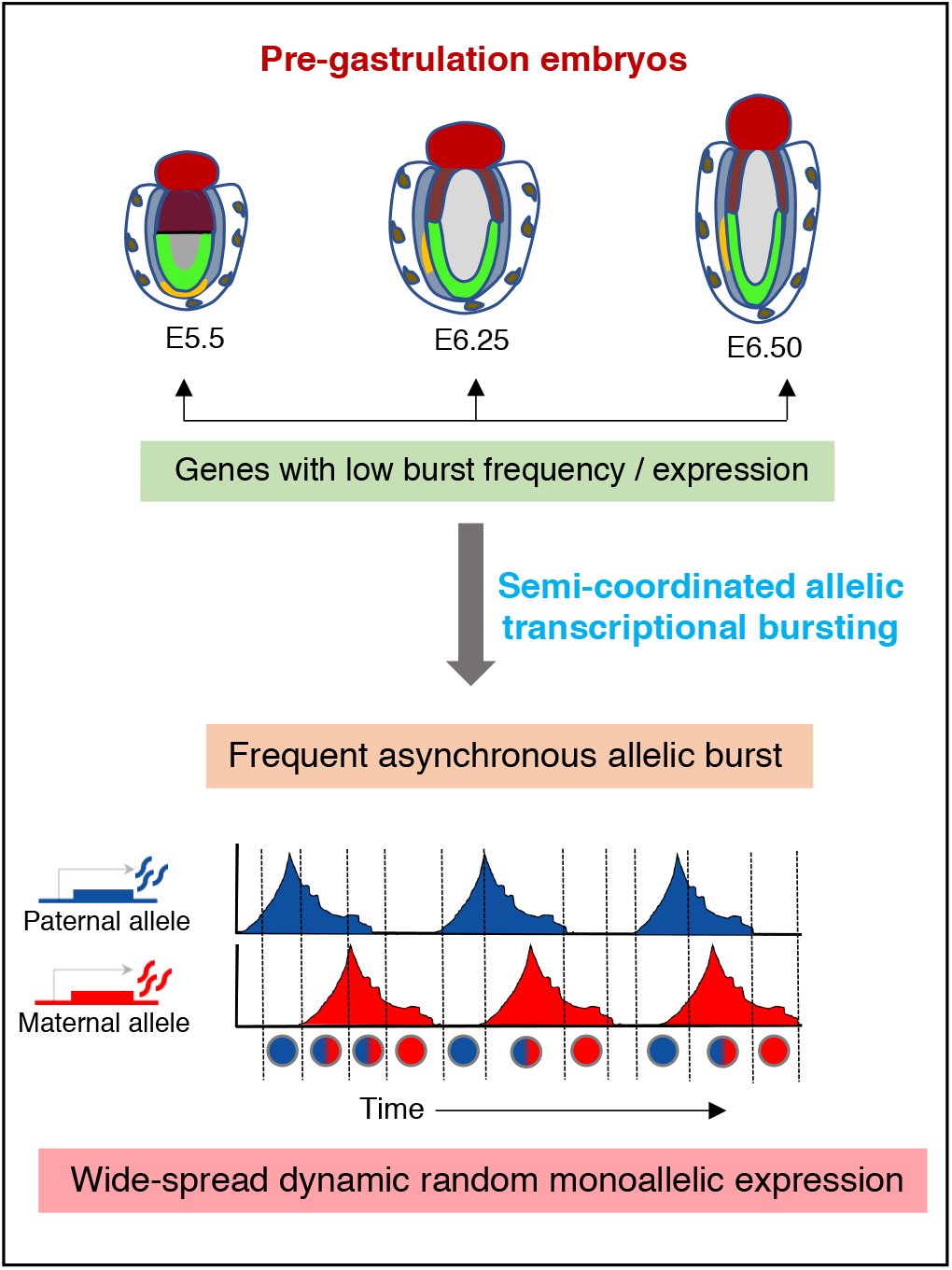

## Introduction

In a diploid eukaryotic cell, both parental alleles of a gene are usually expressed. However, monoallelic expression of genes is common in phenomena like genomic imprinting or X-chromosome inactivation, where a single allele of a gene is expressed (Bartolomei and Ferguson-Smith, 2011; Lyon, 1961; Mandal et al., 2020; Saiba et al., 2018). Surprisingly, recent advances on allele-specific single-cell RNA-seq (scRNA-seq) have revealed that many autosomal genes express monoallelically, which is transient in nature (Deng et al., 2014; Gendrel et al., 2016; Gregg, 2017; Reinius et al., 2016; Reinius and Sandberg, 2015). This wide-spread temporal aRME has been termed as dynamic aRME. Deng et al.’s pioneering study showed that ~12-24% of autosomal genes in a mouse blastomere undergo dynamic RME (Deng et al., 2014). In the same study, analysis of hepatocytes from adult mice and mouse fibroblast cell lines also showed a similar pervasiveness of dynamic aRME (Deng et al., 2014). Subsequently, prevalent dynamic aRME has been reported in various cell-types of mice and humans (Borel et al., 2015; Reinius et al., 2016). However, the prevalence of dynamic aRME during pre-gastrulation development is not known yet. Here, we have profiled the genome-wide pattern of dynamic aRME in different lineages of pre-gastrulation mouse embryos. It is believed that dynamic aRME creates temporal variation among the cells and thereby has the potential to contribute to the cell fate decision, promote cellular plasticity during development. Therefore, profiling the pattern of dynamic aRME during early development is of immense interest.

On the other hand, the origin of dynamic aRME remains poorly understood. It is thought that dynamic aRME is a consequence of stochastic transcriptional burst (Eckersley-Maslin and Spector, 2014; Reinius and Sandberg, 2015). It is known that transcription happens through discrete bursts such that the state of a gene keeps switching randomly from an active to an inactive state, which leads to discontinuous production of mRNA (Raj et al., 2006; Raj and van Oudenaarden, 2008; Suter et al., 2011; Tunnacliffe and Chubb, 2020). For a long time, the transcriptional burst kinetics analysis was mainly based on single-molecule RNA-FISH or live-cell imaging and therefore restricted to a few selected loci of the genome (Raj et al., 2006). Recent advancements in allele-specific expression analysis of many genes at a single cell level have made it possible to analyze transcriptional burst kinetics at allelic level genome-wide more extensively (Ochiai et al., 2020; Sun and Zhang, 2020). However, the kinetics of transcriptional bursting at allelic level remains poorly understood. It is believed that two alleles for most of the genes burst independently (Jiang et al., 2017; Reinius and Sandberg, 2015). Therefore, the abundance of RNA in a cell originating from different alleles can change dramatically over time and lead to dynamic aRME. However, the link between allelic transcriptional burst kinetics and the dynamic aRME has not been explored extensively. In the present study, we have profiled allele-specific transcriptional burst kinetics in different lineages of pre-gastrulation mouse embryos and provided novel insight about the association between allelic burst kinetics and dynamic aRME.

## Results

### Dynamic aRME in different lineages of pre-gastrulation mouse embryos

To investigate the aRME pattern in different lineages of pre-gastrulation mouse embryos, we performed allele-specific gene expression analysis using the available scRNA-seq dataset of E5.5, E6.25, and E6.5 hybrid mouse embryos (Cheng et al., 2019) (Fig. 1A). These embryos are derived from two divergent mouse strains (C57Bl/6J and CAST/EiJ). Therefore, they harbor polymorphic sites between the alleles, which allowed us to perform allelic expression profiles of the genes (Fig. 1A). We segregated the cells into the three lineages: epiblast (EPI), extraembryonic ectoderm (ExE), and visceral endoderm (VE) based on t-distributed stochastic neighbor embedding (t-SNE) analysis and lineage-specific marker gene expression (Fig. S1).

**Figure 1:**
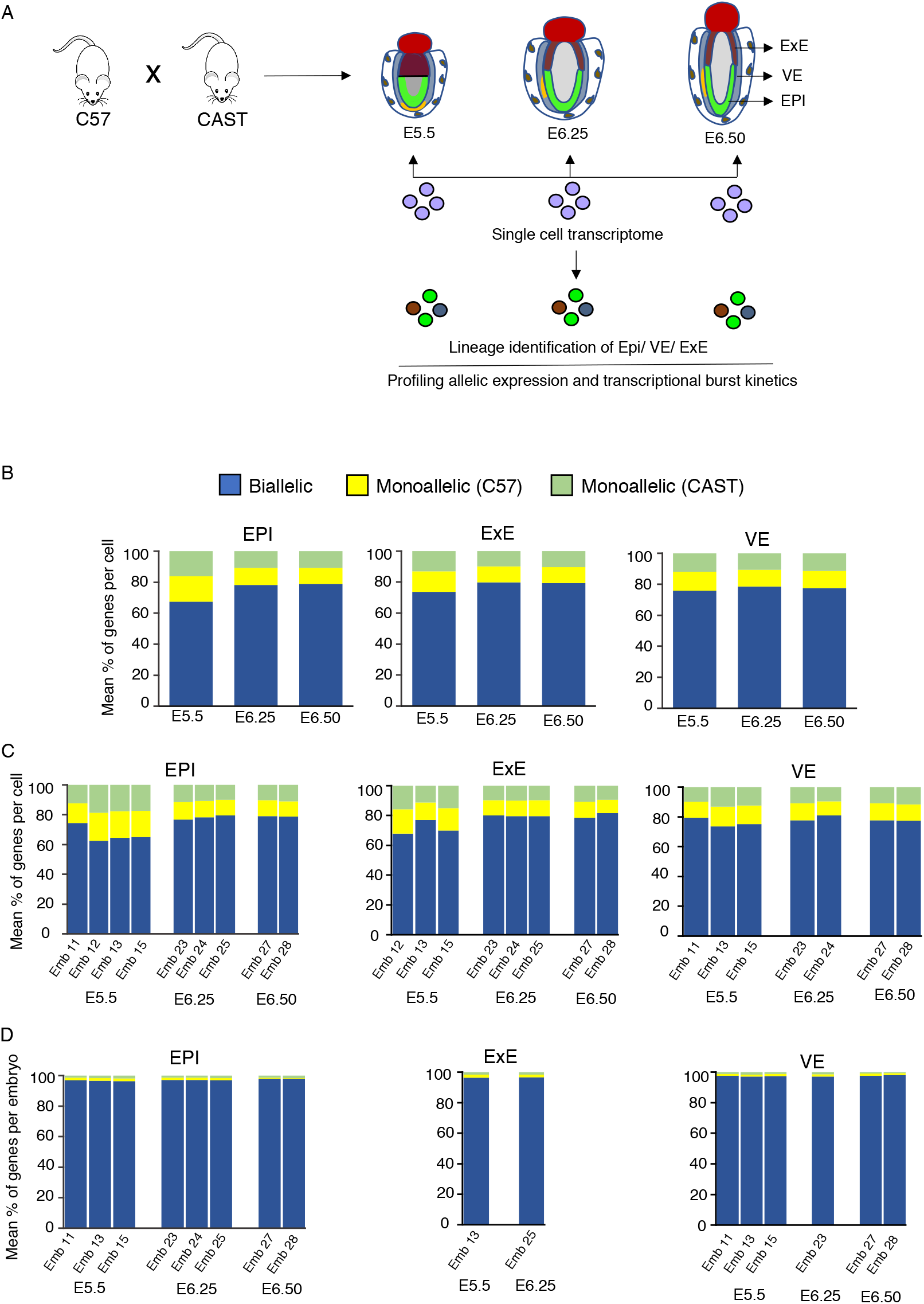
Genome-wide profiling of aRME in different lineages of pre-gastrulation embryos. (A) Graphical outline of the workflow: allelic gene expression and burst kinetics analysis in different lineages (EPI, ExE, and VE) of pre-gastrulation hybrid mouse embryos (E5.5, E6.25, and E6.50) at the single-cell level using published scRNA-seq dataset. Hybrid mouse embryos were obtained from crossing between two divergent mouse strains C57 and CAST. (B) Estimation of mean percent of autosomal genes showing monoallelic expression per cell of each lineage (EPI, ExE, and VE) at different stages (E5.5, E6.25, E6.5). (C) Estimating the mean percent of autosomal genes showing monoallelic expression per cell of each lineage embryo at different stages. (D) Estimating the mean percent of autosomal genes exhibiting monoallelic expression per embryo of each lineage at different stages.

First, we quantified the autosomal genes’ allelic expression pattern in an individual cell of different lineages. We considered a gene as monoallelic if at least 95% of the allelic reads was originated from only one allele. Considering the technical noise can lead to an overestimation of monoallelic expression, especially for lowly expressed genes; we considered those genes with at least mean 10 reads per cell for each lineage of a specific developmental stage. We found with an average of ~15 to 20% of genes showed monoallelic expression either from CAST or C57 allele per cell, and the pattern was almost similar across the three lineages EPI, ExE, and VE of different developmental stages (Fig. 1B). Moreover, each embryo’s allelic expression of different developmental stages showed a very similar pattern (Fig. 1C). Interestingly, per embryo estimation of the mean percent of genes with monoallelic expression by pooling an individual embryo’s cells resulted in significant reduction in fraction of genes with monoallelic expression (0.8-2% genes per embryo) (Fig. 1D). Based on this, we assumed that allelic expression pattern of individual gene might vary cell to cell in each embryo’s lineage at a particular stage. To test our assumption, we investigated the status of the allelic pattern of individual genes across the cells of each lineage of each developmental stage. Indeed, we found a considerable variation of the genes’ allelic status across the cells, indicating the presence of cell-to-cell dynamic aRME (Fig. 2). We observed four different patterns of allelic expression; Cat1: Non-random monoallelic (1 to 2%), Cat 2: random monoallelic with one allele (4-39%), Cat 3: random monoallelic with either allele (30-81%) and Cat 4: biallelic (10 to 29%) (Fig. 2). Altogether, our analysis revealed a high degree of cell-to-cell dynamic aRME (Category 2: 4-39% and Category 3: 30-81%) in each lineage of pre-gastrulation embryos, indicating dynamic allelic expression is a general feature of gene expression affecting many genes during development. We validated our allelic expression analysis through profiling the allelic expression of X-linked genes (Fig. S2).

**Figure 2:**
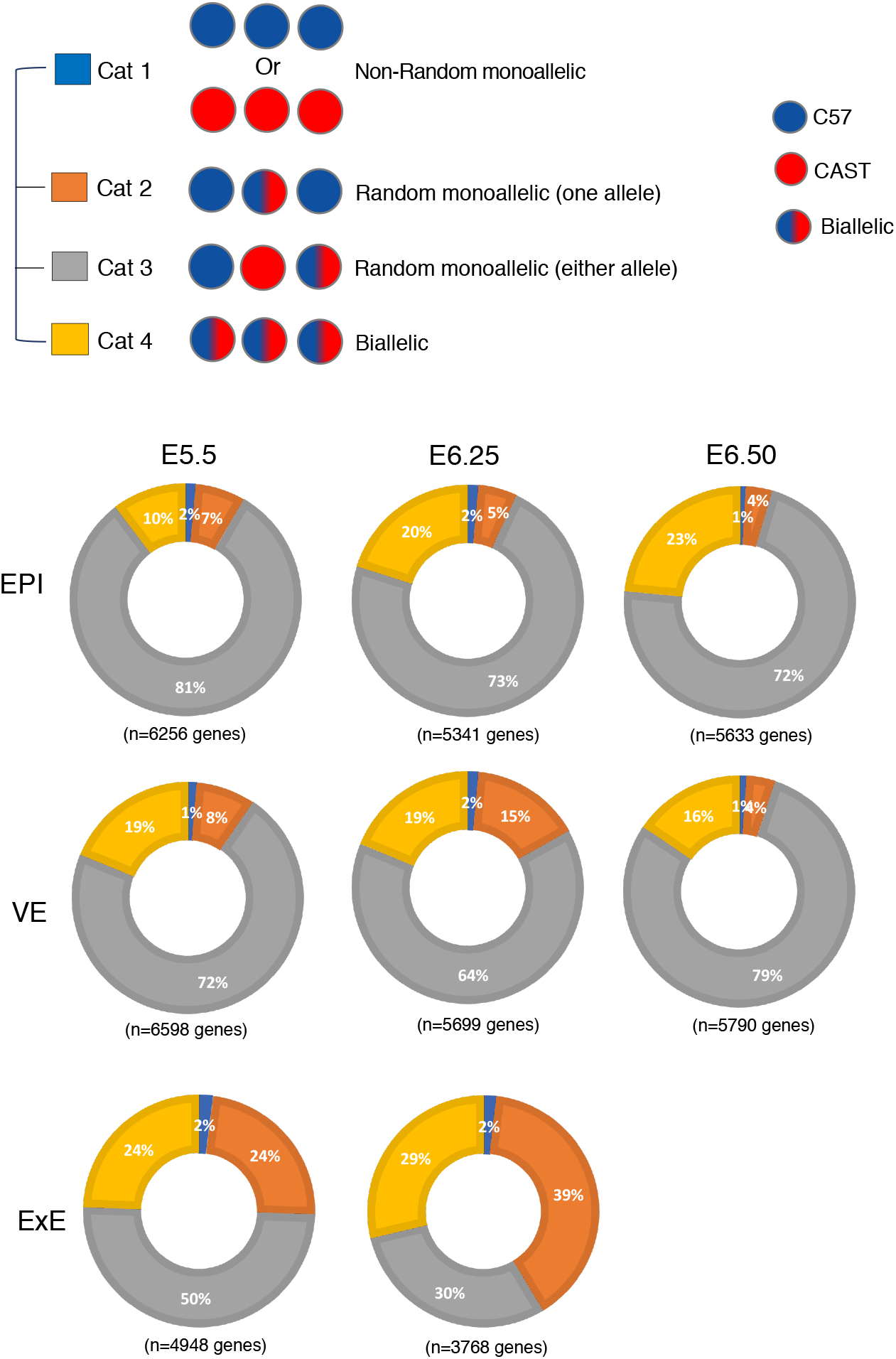
Dynamic aRME in different lineages of pre-gastrulation embryos. Quantification of the percent of genes showing the different category of allelic expression (Cat1: Non-random monoallelic, Cat 2: random monoallelic with one allele, Cat 3: random monoallelic with either allele, Cat 4: biallelic) in three different lineages EPI, ExE, and VE across the various developmental stages.

### Allelic bursting is semi-coordinated

Next, we explored genome-wide allele-specific transcriptional burst kinetics to investigate the link between dynamic aRME and transcriptional bursting. Based on two-state transcription models, transcription occurs in bursts where the state of a gene keeps switching from ON to OFF state (Fig. 3A). Burst kinetics is mainly characterized by burst frequency and burst size. The burst frequency is the rate at which bursts occur per unit time, and burst size is determined by the average number of synthesized mRNA while a gene remains in an active state (Fig. 3A). We used SCALE (Single-Cell Allelic Expression) to determine the genome-wide burst kinetics of autosomal genes in an allele-specific manner (Jiang et al., 2017). Principally, based on the Empirical Bayes framework, SCALE first categorizes the genes to biallelic, monoallelic, and silent using the allele-specific read counts, and then biallelic genes are further classified as biallelic bursty and biallelic non-bursty. Finally, different burst kinetics parameters are deduced for the biallelic bursty genes. We performed burst kinetics analysis for only E6.5 in EPI (n=123 cells) and VE (n=115 cells) cells. For other stages or lineages, there was not a sufficient number of cells for performing SCALE analysis. We considered those autosomal genes (n=5633 genes for EPI and n=5791 genes for VE) for SCALE analysis, which had at least mean ten reads per cell in each lineage. In both E6.5 EPI and VE, we found that most of the genes (70-82%) showed bursty expression (Fig. 3B; Supplementary file 1). Next, we compared the burst kinetics between the alleles of biallelic bursty genes. Interestingly, we found that the alleles of most of the genes showed similar burst kinetics, i.e., they had identical burst frequency and size (Fig. 3C & 3D, Supplementary file 1). Only 48 out of 3861 bursty genes (EPI) and 90 out of 4705 bursty genes (VE) showed significantly different allelic burst frequency after FDR correction (Fig. 3C). On the other hand, very few genes showed significantly different allelic burst size (Fig. 3D). Next, we determined the independence of allelic transcriptional burst. We plotted the percent of cells expressing neither allele (p_0_) with the percent of cells expressing both alleles (p_2_), as depicted in Fig. 3E. In the perfect independent model, most of the genes (black dots) should lie across the red curve, whereas perfect coordination model genes should lie near the diagonal blue line. Interestingly, we found that most of the genes reside in the middle of between the red and diagonal blue lines, indicating that allelic bursting is neither entirely independent nor perfectly coordinated (Fig. 3E). Additionally, the null hypothesis of independence was rejected for most of the genes (Supplementary file 1). Altogether these results suggested that alleles of most of the genes have similar burst kinetics; however, allelic bursting was neither entirely independent nor perfectly coordinated.

**Figure 3:**
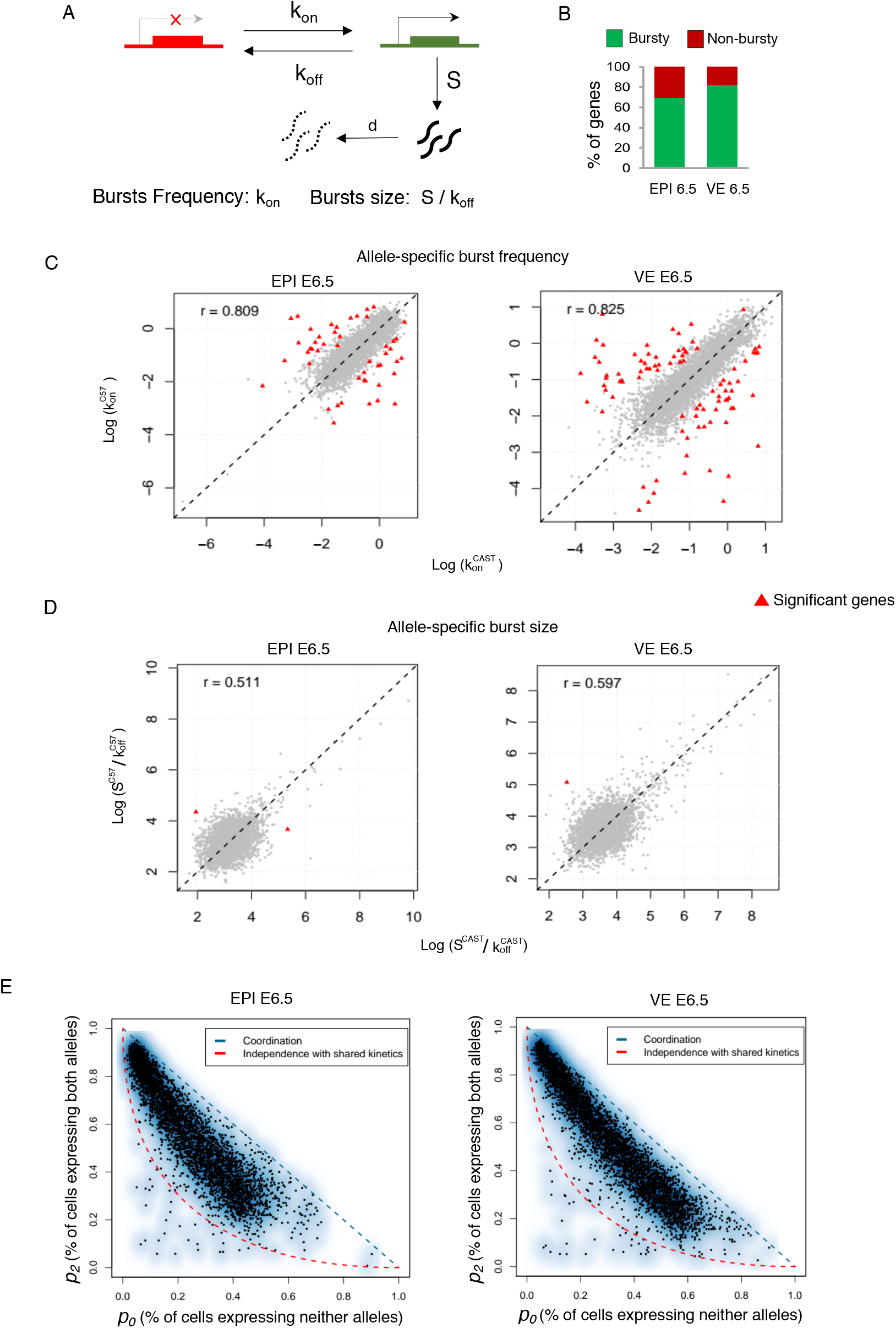
Genome-wide profiling of transcriptional burst kinetics. (A) Representation of the two-state model of transcription. k_on_: the rate at which a gene becomes transcriptionally active (from off to on); k_off_: the rate at which a gene becomes inactive (from on to off); S is the rate of transcription, while the gene is active; d is the rate of mRNA decay. Burst kinetics is characterized through burst frequency (k_on_) and the burst size (S/k_off_). (B) Estimating the proportion of autosomal genes with bursty expression in EPI and VE cells of the E6.5 stage. (C) The burst frequency of the two alleles of most of the genes was highly similar in EPI and VE cells of the E6.5 stage (r= 0.80 and 0.82 respectively). Genes having significantly different allelic burst frequency marked as a red triangle. (D) A similar burst size of the two alleles of most of the genes in EPI and VE cells of the E6.5 stage (r= 0.51 and 0.59, respectively). Genes having significantly different allelic burst sizes are marked as a red triangle. (E) Smooth scatter plot showing independence of allelic, transcriptional bursting in EPI, and VE cells of E6.5 stage. p_0_ is _the_ percent of cells expressing neither allele. p_2_ is the percent of cells expressing both alleles. Black points representing individual genes. The diagonal blue line (p_0_ + p_2_ = 1) represents coordinated bursting, whereas the red curve represents perfect independent bursting with shared kinetics.

Next, to get a quantitative assessment of the dependence between the alleles, we constructed a simple two-component stochastic model, inspired from the classic two-state model of transcription (Peccoud and Ycart, 1995) (Fig. S3). This model describes two alleles identical kinetic parameters in terms of their probabilities to switch ‘on’ and ‘off’. To enable bursting, we introduce two more parameters – stayOn and stayOff - which represent the change in probability of an allele staying in its present state. To model the dependency of the transcription of one allele on the other, we assumed that the probability of an allele turning and staying on is multiplied by a parameter lambda, when the other allele is on. Therefore, lambda>1 describes that an allele in ‘on’ state facilitates the other allele also being ‘on’ (the higher the value of lambda, the higher the effect), lambda =1 describes independent bursting of the two alleles, and lambda < 1 describes that an allele being ‘on’ disfavours the other allele staying ‘on’. In total for both the alleles, we have 5 parameters in the model. The on and off probabilities are chosen between 0 and 1 and lambda values are chosen between 0.01 and 100. First, we validated our model through simulations for independently expressing alleles through testing it against the theoretical relationship between probability of both alleles being off (p0) and both alleles being on (p2). Given the alleles both have the same on probability (p), p0 is given as (1-p)*(1-p) and p2 is given as p*p. Therefore, p2 and p0 are related via the following equation:

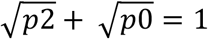

We found that the values obtained via numerical simulations (Fig 4A) lie very close to this curve and distributed symmetrically along the curve (Fig 4A inset), thus validating the model. Next, to understand how various parameters affect the placement of genes in the p0-p2 plot, we divided the data obtained from simulations into four regions (Fig 4B). The yellow region represents points below the independence curve, orange dots are those lying on the absolute dependence curve, black dots denote the region between the two curves, and red dots are for the region with high p0 (which is not seen in experimental data). Among the four, the experimental data points align best with the black region in the plot. The original distribution of different parameters for all simulations is shown in Fig 4C. First, we looked at the region with high p0. The distribution of parameters revealed that these points had very low on probability and high values of lambda (Fig 4D). A scatter plot between these two parameters further shows that for high values of lambda, the probability to switch to an ‘on’ state must be very small (Fig 4D inset). In the experimental data, the fact that none of the points lie in this region may possibly indicate that the data collection method is not sensitive to genes with on probabilities lower than 0.1. Next, we looked at the points below the independence curve (Fig 4E). The lambda values for these points do not look characteristically different from those seen in the cumulative distribution (i.e. all points taken together). These points correspond to lambda values below 1 and relatively lower off rates. Conversely, the points on the perfect dependence line have high lambda values and a lower peak for the off probability, suggesting that very high lambda is needed for absolute dependence (Fig 4F). For the black region, lambda values lie largely between 10-50, which happens to be the middle region of all lambda values sampled (Fig 4G). The distributions of other parameters do not have observable differences from the cumulative distributions (Fig 4C). Together, we concluded that the experimentally observed data points correspond to cases with probability of turning on both alleles higher than 0.1 and have moderate levels of the co-dependence parameter lambda. Next, to establish a causal connection, we applied these conditions on our parameter sampling, and numerical simulations showed a distribution of genes in the p0-p2 axis very similar to the experimental observations (Fig 4H). Specifically, all genes lie between the theoretical curves of independence and absolute dependence, with no genes having p0 greater than 0.6. Put together, this quantitative analysis highlights the mechanistic underpinnings underlying observed experimental data.

**Figure 4:**
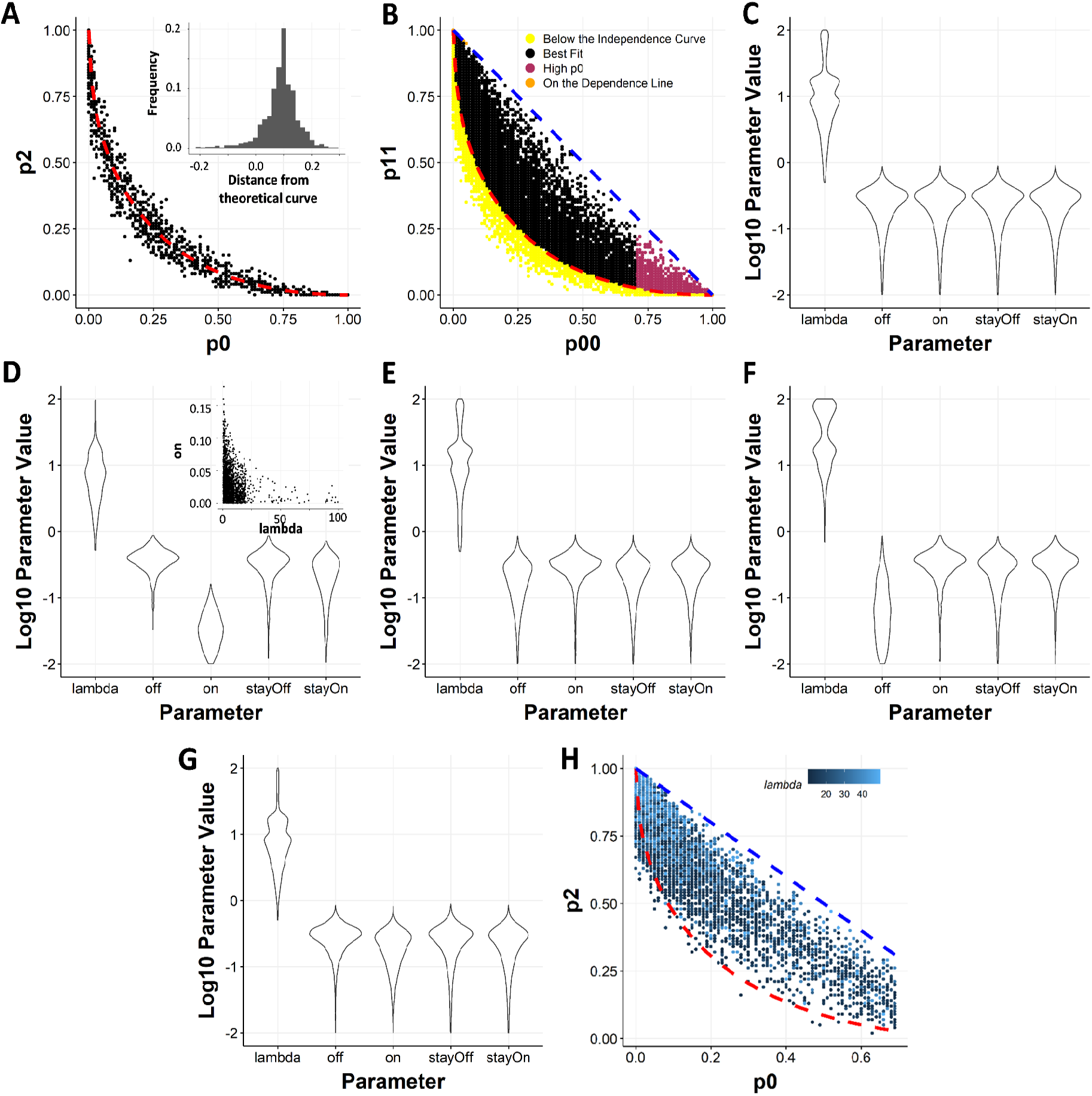
Simulation of the dependence between allelic bursting. (A) Comparison of simulation results for independent allele bursting with theoretical curve representing independent bursting on the p0-p2 plot. A distribution of distance of the points from the curve is shown in the inset. (B) Classification of simulation data based on their position in the p0-p2 plot with respect to the theoretical curves. (C) Violin plots showing distributions from which each of the 5 parameters were sampled. (D) Parameter distribution for the genes represented with maroon dots (p0>0.7) in B. Inset shows a scatterplot between lambda and on probability. (E) Parameter distribution for the genes represented with yellow dots (below the theoretical independence curve) in B. (F) Parameter distribution for the genes represented with orange dots (strong dependence) in B. (G) Parameter distribution for the genes represented with black dots (genes lying between the two theoretical curves) in B. (H) Scatterplot between p0 and p2 constructed by using two constraints on the parameter values: on > 0.1 and 10<lambda<50.

### Dynamic aRME is linked to allele-specific transcriptional burst kinetics

Next, we delineated the correlation between allelic transcriptional burst kinetics and dynamic aRME. First, we wanted to see if there any correlation between bursty gene expression and dynamic aRME. Interestingly, we found that most dynamic aRME genes (Cat 2 & Cat 3) showed bursty expression (Fig. 5A). Especially for Cat 3 aRME genes, more than 92% of genes showed bursty expression (Fig. 5A). On the other hand, most biallelic genes (Cat 4) for EPI cells showed non-bursty expression (Fig. 5A). Altogether, these results suggested that dynamic aRME is generally linked with bursty expression. Next, we examined if there any correlation between the allelic expression of genes with the allelic burst kinetics. To test this, we performed a pairwise correlation test between different burst kinetics parameters, and the sum of allelic read counts for each gene across the cells (Fig. 5B). We found that the total expression of alleles is positively correlated (r=0.65-0.77) with allelic burst frequency. On the other hand, although allelic expression was positively correlated with the burst size (r=0.12-0.18) and the proportion of unit time, the allele remains active (r=0.23-0.34), the correlation value was much lower compared to the burst frequency. To get more insight into this aspect, we compared the burst frequency and burst size of alleles with the percent of cells expressing that corresponding allele. Interestingly, we found that the proportion of cells express one allele of genes is dependent on the burst frequency of that allele rather than burst size (Fig. 5C). Overall, we found that the proportion of cells expressing one allele of genes increases parallel way with the increase in burst frequency. Similarly, we compared the mean expression of alleles with the allelic burst frequency and burst size and found that mean expression level substantially dependent on allelic burst frequency instead of burst size (Fig. 5D). Allelic expression was directly proportional to the allelic burst frequency such that alleles expressing high showed high allelic burst frequency, low had low allelic burst frequency. Altogether, these analyses suggested that burst frequency among the different kinetics parameters is crucial for monoallelic gene expression. Next, we delineated if dynamic aRME is dependent on the overall expression level. Interestingly, comparison of expression level between bursty vs. non-bursty genes revealed that non-bursty genes always have significantly higher expression than the bursty genes (Fig. 6A). Next, we hypothesized that the proportion of cells with the monoallelic expression might depend on the gene’s expression level. We analyzed the correlation between gene expression level and percent of cells showing the monoallelic expression for that gene to test our hypothesis. As expected, we found a high negative correlation (r= −0.58 to −0.61) (Fig. 6B). Altogether, these results indicated that the extent of a gene’s monoallelic expression depends on its expression level and allelic burst frequency. Our observation and analysis proposed a model highlighting how transcriptional burst kinetics can contribute to the dynamic aRME (Fig. 6C). We propose that bursty genes with asynchronous allelic burst kinetics build up the dynamic aRME landscape. Genes with lower expression and/or lower burst frequency frequently undergo monoallelic expression (Fig. 6C). On the other hand, genes with high expression and/or high allelic burst frequency express most of the time (Fig. 6C) biallelically.

**Figure 5:**
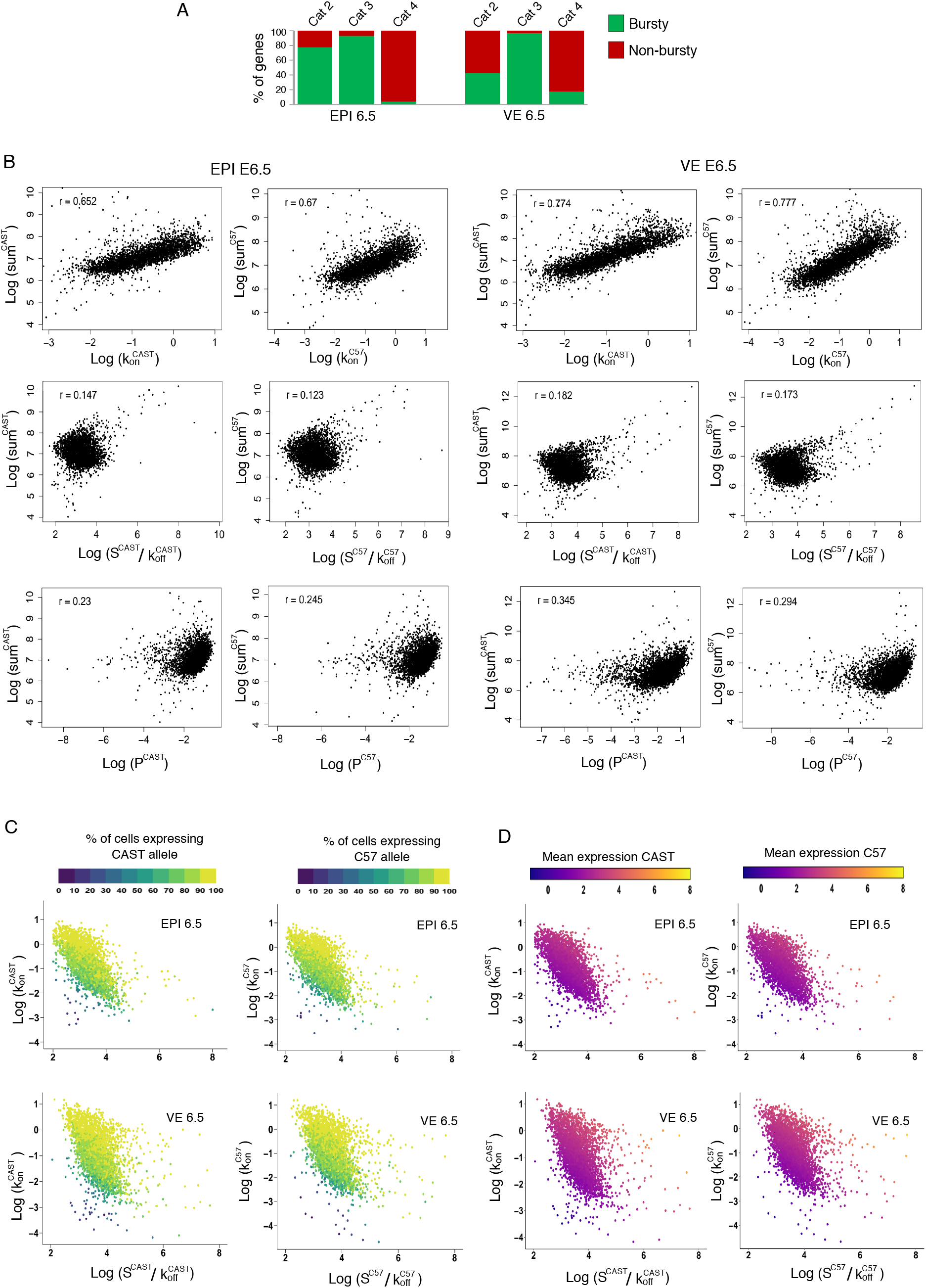
Association between burst kinetics and dynamic aRME. (A) Quantification of **the** proportion of dynamic aRME genes (Cat 2 & Cat 3) with bursty expression and proportion of biallelic genes (Cat 4) having bursty expression in EPI and VE cells of E6.5 stage. (B) Pairwise correlation between different allelic bursting kinetics parameters (burst frequency: k_on_^CAST^ and k_on_^C57^; Proportion of unit time that the gene stays in active form: *p*^CAST^ = k_on_^CAST^/(k_on_^CAST^ + k_off_^CAST^); *p*^C57^ = k_on_^C57^/(k_on_^C57^ + k_off_^57^); burst size: S^CAST^/k_off_^CAST^ and S^C57^/k_off_^C57^) and expression level (sum of normalized allelic read counts (log)) of the alleles in EPI and VE cells of E6.5 stage. (C) Scatter plot representing an estimate of burst size and a burst frequency of the CAST and C57 allele of autosomal genes. The gene’s color is profiled based on the percent of cells expressing the CAST or C57 allele. (D) Scatter plot representing an estimate of burst size and burst frequency of the CAST and C57 allele of autosomal genes. The color of the gene is depicted based on the mean allelic expression.

**Figure 6:**
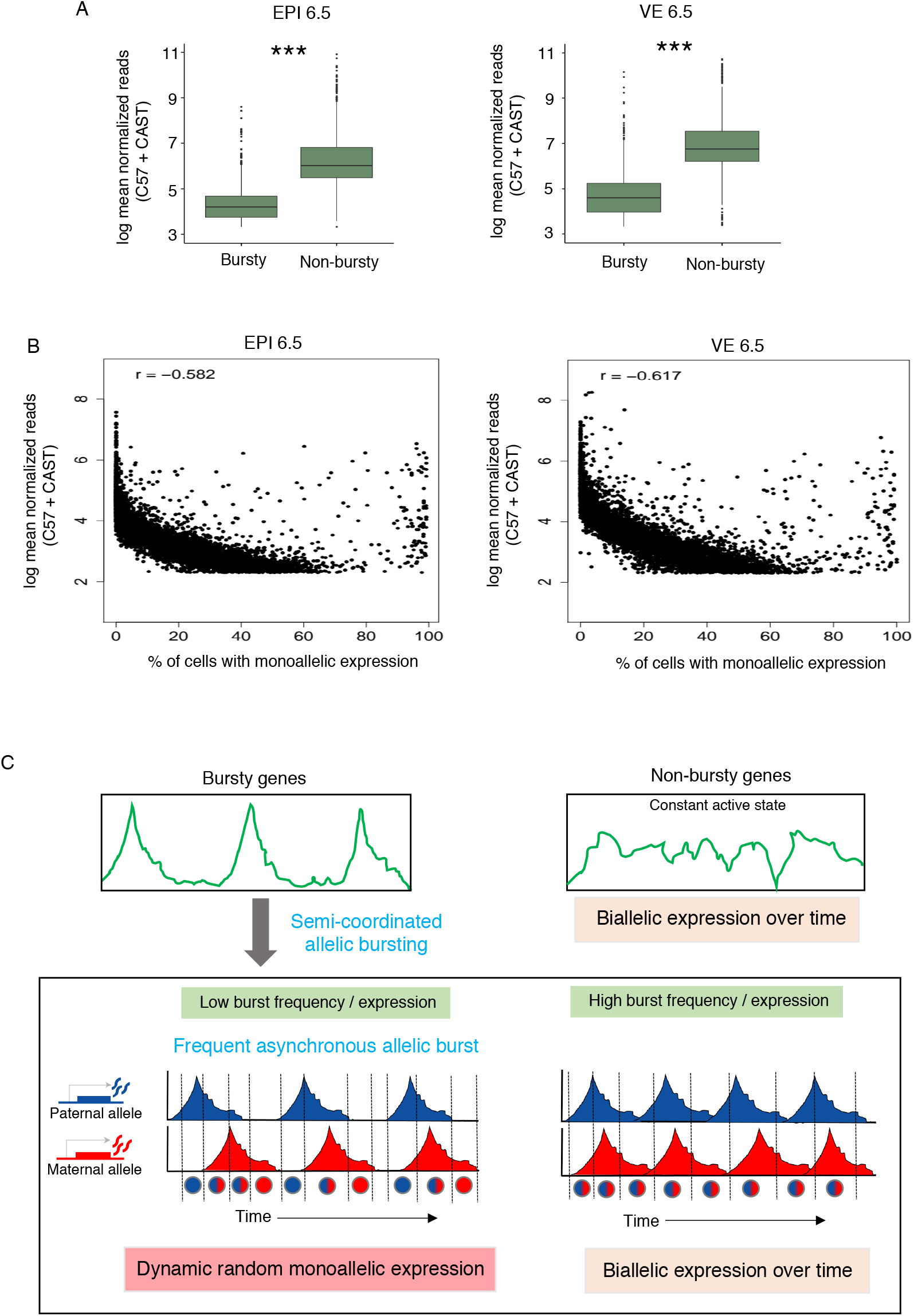
Relation of bursting and dynamic aRME with the gene expression level. (A) Comparison of the expression level of bursty vs. non-bursty genes in EPI and VE cells of the E6.5 stage. (B) Correlation plot between the mean expression of gene and percent of cells showing monoallelic expression for that gene (r= −0.58 in EPI and −0.61 in VE). (C) Model representing how transcriptional burst kinetics can lead to dynamic aRME.

## Discussion

It is believed that dynamic aRME creates temporal variation among the cells and thereby can contribute to the cell fate decision and promote cellular plasticity during development (Gregg, 2017; Huang et al., 2018; Montag et al., 2018; Ng et al., 2018). Therefore, investigating dynamic aRME during early development is crucial. In the present study, we show wide-spread dynamic aRME in different lineages of pre-gastrulation mouse embryos. Notably, dynamic aRME is more prevalent (~69-88% genes) in pre-gastrulation embryos in comparison to the blastomeres (~12-24%) reported by Deng et al. (Deng et al., 2014) (Fig. 2). This robust increase in the fraction of dynamic aRME in pre-gastrulation embryos indicates that dynamic allelic expression is a general feature of gene expression affecting many genes during development. Separately, among these aRME genes, some gene’s allelic expression pattern might be mitotically heritable, as reported earlier (Eckersley-Maslin et al., 2014; Gendrel et al., 2014; Gimelbrant et al., 2007; Jeffries et al., 2016; Zwemer et al., 2012). In the future, investigation on the clonal cell population can disentangle the mitotically stable aRME from the dynamic aRME.

On the other hand, we have profiled genome-wide allele specific burst kinetics of autosomal genes to understand the implication of allelic bursting on the dynamic aRME. We found that majority of the autosomal genes have bursty expression, and alleles of most of the genes have similar burst kinetics, which is consistent with previous reports in other cell types (Fig. 3B, C &D) (Jiang et al., 2017). However, we found that allelic bursting is not perfectly independent instead, it happens in a semi-coordinated fashion (Fig. 3E & Fig. 4). Finally, we demonstrate that dynamic aRME is linked to semi-coordinated allelic bursting. We show that majority of dynamic aRME genes have bursty expression, whereas most of the biallelic genes were found to be non-bursty. Moreover, we found that extent of dynamic aRME is determined by burst frequency rather than burst size or how long an allele remains active. Notably, we found that dynamic aRME was highly dependent on the overall expression level of a gene. Altogether, we propose that semi-coordinated allelic bursting for the genes with lower burst frequency leads to frequent asynchronous allelic bursting, thereby creating wide-spread dynamic aRME (Fig. 6C). On the other hand, non-bursty genes or bursty genes with high allelic burst frequency and/or high expression level exhibit frequent biallelic expression (Fig. 6C). In the future, more investigations are necessary to determine whether dynamic aRME is a mere consequence of stochastic transcriptional bursting or they have a wide range of biological significances such as in development and disease. Additionally, extensive studies are necessary to understand the regulatory network behind semi-coordinated allelic bursting. In perfect independent model regulation of allelic expression should be autonomous, whereas, in an alternative model of perfect dependence, there can be shared allelic expression regulation. We believe that autonomous as well as shared regulation of the alleles results in semi-coordinated transcriptional bursting. Interestingly, a recent study has shown that chromatin conformations are variable between alleles, and each allele can behave independently, indicating that regulation of allelic expression can be self-governing (Finn et al., 2019). Moreover, it has been shown that while allelic burst frequency is regulated through enhancer, burst size is controlled by the core promoter (Larsson et al., 2019). Moreover, a recent report suggests stochastic switching between methylated to unmethylated state at many regulatory loci occurs in a sequence-dependent manner, which can be another mechanism behind stochastic transcriptional bursting (Onuchic et al., 2018).

## Methods

### Data acquisition

Single-cell transcriptome datasets used for this study were acquired from Gene Expression Omnibus (GEO) under the accession number GSE109071 (Cheng et al., 2019). For our research, we analyzed a single-cell dataset generated from E5.5, E6.25, and E6.50 hybrid mouse embryos (C57BL/6J × CAST/EiJ). E5.5 and E6.25 embryos were derived from the following cross: C57(F) × CAST(M), whereas E6.5 were derived from CAST(F) × C57(M).

### Lineage identification

All the single cells (510 cells) of different stages were subjected to a dimension reduction algorithm using t-distributed stochastic neighbor embedding (t-SNE) to identify lineages. Three thousand most variable genes were used for the analysis. t-SNE was performed using Seurat (version 3.1.5) (Butler et al., 2018; Stuart et al., 2019). The allocation of each cluster to cell lineages to EPI, ExE, and VE lineages was based on the expression of bona fide marker genes: Oct4 for EPI, Bmp4 for ExE, and Amn for VE.

### Allele-Specific Expression and burst kinetics analysis

For allelic expression analysis of genes, first, we constructed *in silico* CAST specific parental genome by incorporating CAST/EiJ specific SNPs into the GRCm38 (mm10) reference genome using VCF tools (Danecek et al., 2011). CAST specific SNPs were obtained from the Mouse Genomes Project (https://www.sanger.ac.uk/science/data/mouse-genomes-project). Reads were mapped onto both C57BL/6J (mm10) reference genome and CAST/EiJ *in silico* parental genome using STAR with no multi-mapped reads. To exclude any false positive, we only considered those genes with at least 1 informative SNPs (at least 3 reads per SNP site). In genes having more than 1 SNP, we took an average of SNP-wise reads to have the allelic read counts. We normalized allelic read counts using spike-in. We considered those genes which had at least mean 10 reads per cell for each lineage of a specific developmental stage. Allelic expression was calculated individually for each gene using formula = (Maternal/Paternal reads) ÷ (Maternal reads + Paternal reads). A gene was considered monoallelic if at least 95% of the allelic reads came from only one allele. We performed allele-specific burst kinetics analysis using SCALE (Jiang et al., 2017).

## Author’s Contribution

SG conceptualized and supervised the study. Bioinformatic analyses was performed by HCN. DC and SM helped with the analysis. MKJ and KH performed Simulation. SG, HCN, MKJ and KH wrote the manuscript. Final manuscript was approved by all the authors.

## Acknowledgments

We thank R.V. Pavithra for her help in artwork. Study is supported by DBT grant (BT/PR30399/BRB/10/1746/2018), DST-SERB (CRG/2019/003067), DBT-Ramalingaswamy fellowship (BT/RLF/Re-entry/05/2016) and Infosys Young Investigator award to SG. We also thank DST-FIST [SR/FST/LS11-036/2014(C)], UGC-SAP [F.4.13/2018/DRS-III (SAP-II)] and DBT-IISc Partnership Program Phase-II (BT/PR27952-INF/22/212/2018) for infrastructure and financial support.

---

**Figure S1:**
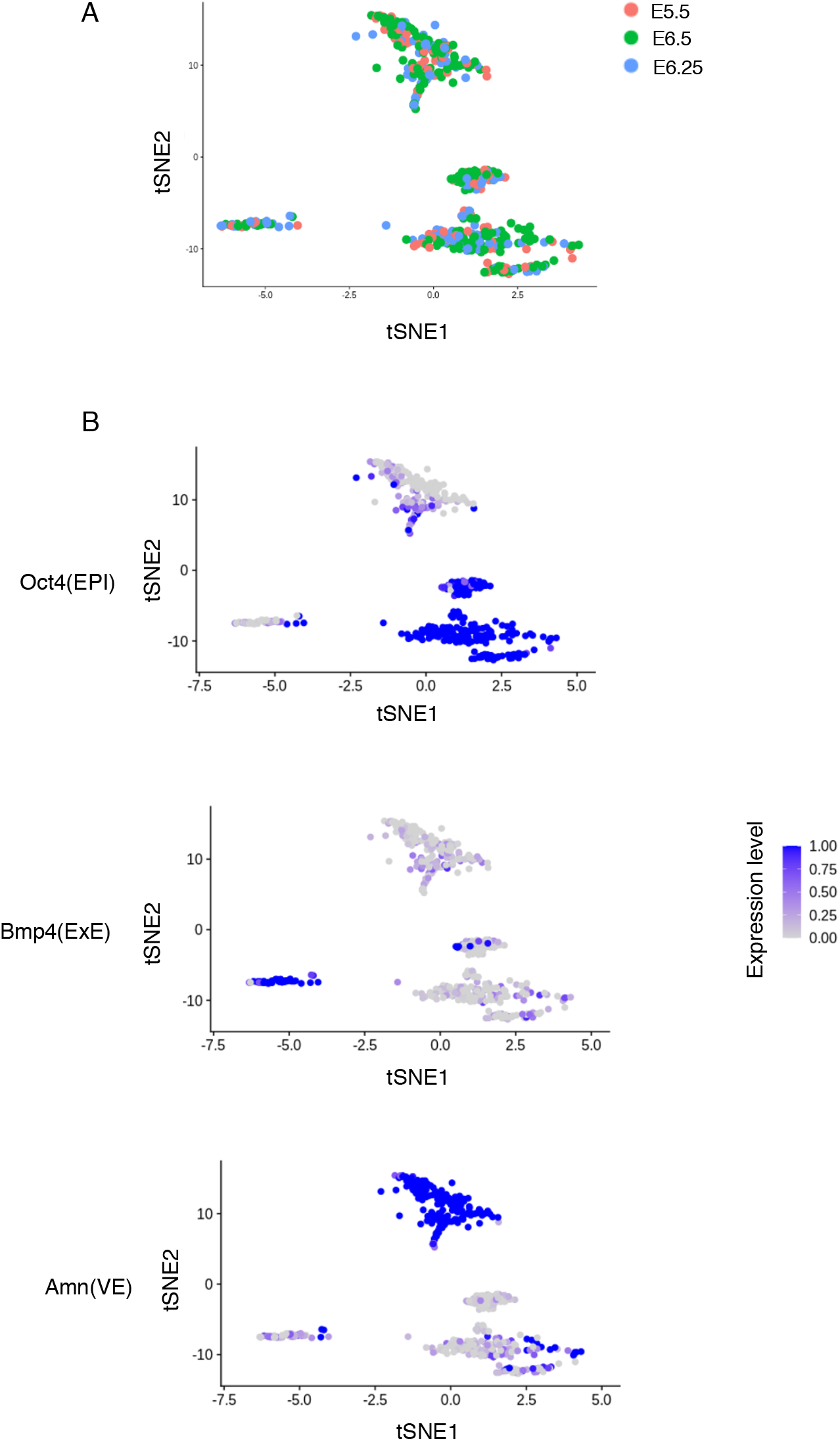
Lineage profiling of pre-gastrulation mouse embryos based on single cell transcriptomics. (A) Clustering of all cells (n=510) from the three different stages (E5.5, E6.25 and E6.50) into two principal dimensions using t-SNE analysis based on 3000 most variable genes. (B) Representation of lineage specific marker expression of the clustered cells generated in t-SNE plot: Pou5f1 for EPI, Bmp4 for ExE and Amn for VE.

**Figure S2:**
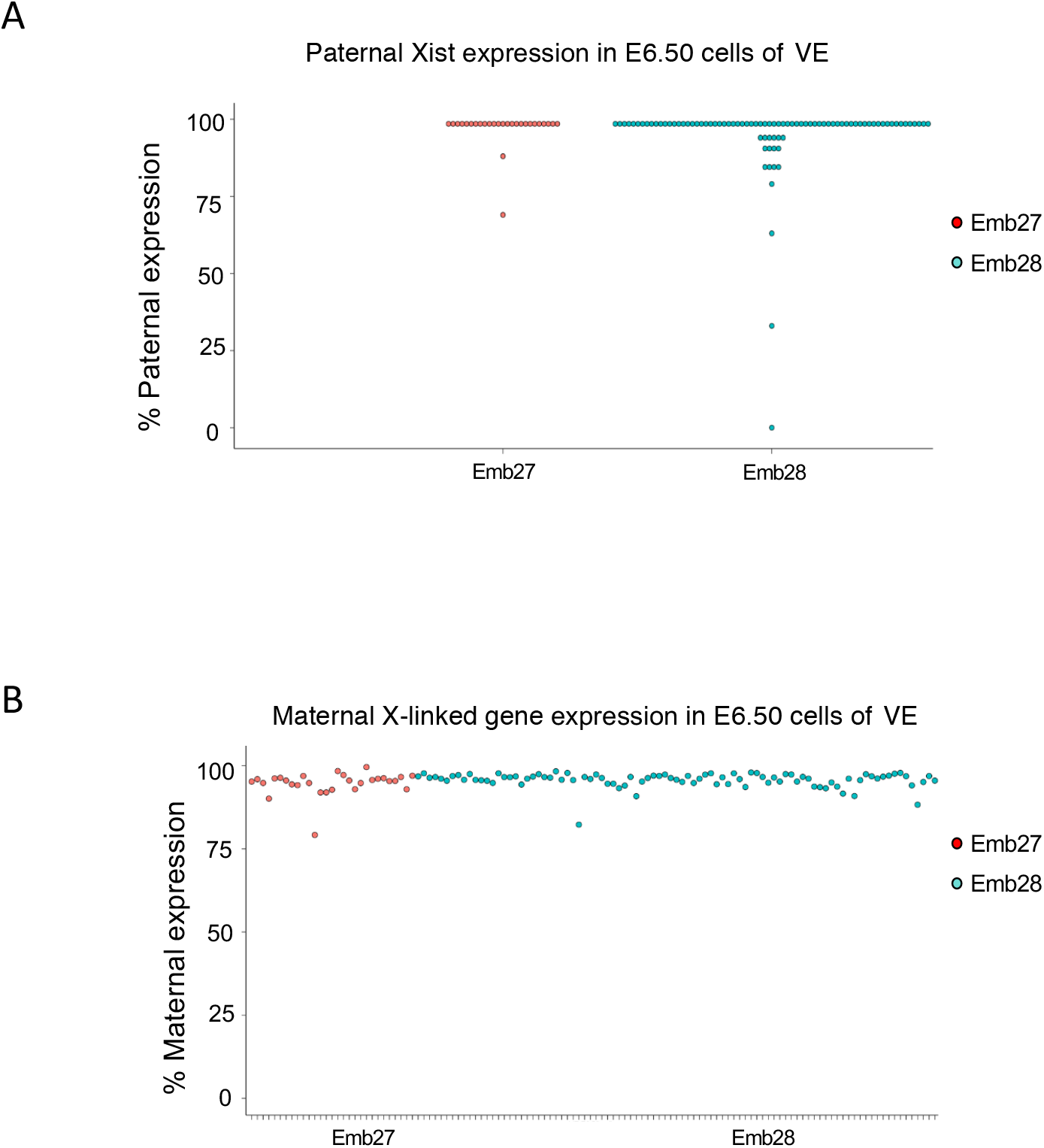
Expression of *XIST* from paternal X-chromosome and X-linked genes from maternal X-chromosome in E6.5 female VE cells. Female VE cells undergo imprinted X-inactivation and therefore paternal X-chromosome is chosen as the inactive-X chromosome. *XIST* long noncoding RNA exclusively expresses from the inactive-X chromosome. (A) As expected, we found in almost all cells except few, allelic expression of *XIST* originated from paternal-X chromosome. (B) Profiling allelic expression of X-linked genes from maternal allele, showed >90% of expression from the active maternal-X chromosome almost in all cells and thus validating the accuracy of the allelic expression analysis method.

**Figure S3:**
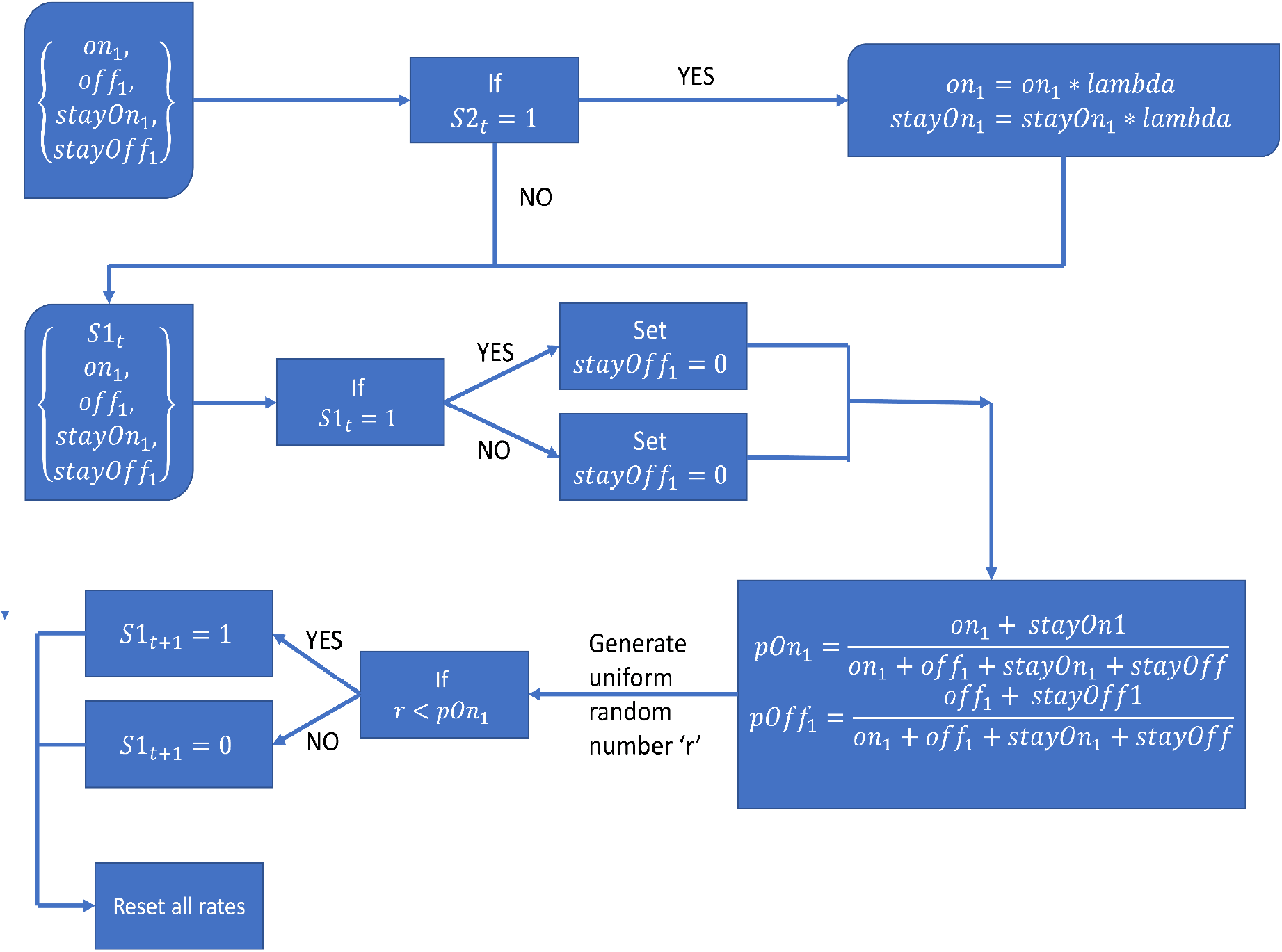
The model describes a gene with two alleles, with identical parameters. The simulation algorithm is described in the flow chart for one allele. The allele is described by the parameters {on1, off1, stayOn1, stayOff1}. At the beginning of the simulation, each of these parameters are sampled from a uniform distribution ranging from 0 to 1. The on/off state of the alleles at any given time t is given by {S1t, S2t}. The dependence parameter lambda is sampled from a uniform distribution ranging from 0.01 to 100.

## Notes

### Competing Interest Statement

The authors have declared no competing interest.

